# Spatial Control of Neuronal Metabolism Through Glucose-Mediated Mitochondrial Transport Regulation

**DOI:** 10.1101/372284

**Authors:** Anamika Agrawal, Gulcin Pekkurnaz, Elena F. Koslover

## Abstract

Eukaryotic cells modulate their metabolism by organizing metabolic components in response to varying nutrient availability and energy demands. In the axons of mammalian neurons, mitochondria have been shown to respond to glucose levels by halting active transport preferentially in high glucose regions. Here, we employ quantitative modeling to explore the physical limits on spatial organization of organelles through such regulated stopping of processive motion, as well as the consequences to cellular metabolism. We delineate the role of key parameters, including cellular glucose uptake and consumption rates, that are expected to modulate mitochondrial distribution and metabolic response in spatially varying glucose conditions. Our quantitative estimates indicate that physiological brain glucose levels fall within the limited range necessary for metabolic enhancement, making this a plausible regulatory mechanism for neuronal metabolic flexibility in the presence of spatially heterogeneous glucose. These findings highlight the role of spatial organization in the regulation of neuronal metabolism, while providing a quantitative framework for the establishment of such organization by control of organelle trafficking.

## INTRODUCTION

Cellular metabolism comprises an intricate system of reactions whose fine-tuned control is critical to cell health and function. A number of quantitative studies have focused on metabolic control through modulating reactant and enzyme concentrations and turnover rates [1, 2]. However, these studies generally neglect the spatial organization of metabolic components within the cell. By localizing specific enzymes in regions of high metabolic demand[3, 4], as well as clustering together consecutively acting enzymes[5], cells have the potential to substantially enhance their metabolism.

Spatial organization is particularly critical in highly extended cells, such as mammalian neurons, whose axons can grow to lengths on the meter scale. Metabolic demand in neurons is spatially and temporally heterogeneous, with especially rapid ATP turnover found in the presynaptic boutons[6], and ATP requirements peaking during synaptic activity and neuronal firing[7–9]. Neurons rely primarily on glucose as the energy source for meeting these metabolic demands[10]. Due to the long lengths of neural processes, the glucose supply can vary substantially over different regions of the cell[8, 9, 11]. In myelinated neurons, for instance, it has been speculated that glucose transport into the cell is localized primarily to narrow regions around the nodes of Ranvier[12–14], which can be spaced hundreds of microns apart[15, 16]. Glucose transporters in neurons have also been shown to dynamically mobilize to active synapses, providing a source of intracellular glucose heterogeneity[17]. Furthermore, in the mammalian brain, extracellular glucose levels vary substantially between different brain regions, resulting in spatially heterogeneous nutrient access[18]. Individual axons have been shown to span across multiple regions of the brain[19], enabling them to encounter regions with varying glucose concentrations.

Most ATP production in neurons occurs within mitochondria: motile organelles that range from interconnected networks to individual globular structures that extend throughout the cell. As energy powerhouses and metabolic signaling centers of the cell, mitochondria are critical for neuronal health [20]. Their spatial organization within the neuron plays a pivotal role in growth and cell physiology [21]. Defects in mitochondrial transport are involved in the pathologies of several neural disorders such as peripheral neuropathy and Charcot-Marie-Tooth disease [22, 23].

A number of studies have shown that mitochondria are localized preferentially to regions of high metabolic demand, such as the synaptic terminals [21, 24]. Such localization can occur via several molecular mechanisms, mediated by the Miro-Milton mitochondrial motor adaptor complex that links mitochondria to the molecular motors responsible for transport[25]. Increased Ca^2+^ levels at active synapses lead to loading of calcium binding sites on Miro, releasing mitochondria from the microtubule and thereby halting transport[26, 27]. High glucose levels can also lead to stalling, through the glycosylation of motor adaptor protein Milton by the glucose-activated enzyme *O*-GlcNAc transferase (OGT)[28]. This mechanism has been shown to lead to mitochondrial accumulation at glucose-rich regions in cultured neurons[28]. It is postulated to regulate mitochondrial spatial distribution, allowing efficient metabolic response to heterogeneous glucose availability.

Mitochondrial positioning relies on an interplay between heterogeneously distributed diffusive signaling molecules (such as Ca^2+^ and glucose), their consumption through metabolic and other pathways, and their effect on motor transport kinetics. While the biochemical mechanisms and physiological consequences of mitochondrial localization have been a topic of much interest in recent years[25, 29], no quantitative framework for this phenomenon has yet been developed.

In this work we focus on glucose-mediated regulation of mitochondrial transport, developing quantitative models to examine the consequences of this phenomenon for metabolism under spatially varying glucose conditions. Our approach relies on a reaction-diffusion formalism, which describes the behavior of species subject to both consumption and diffusion [30]. Reaction-diffusion systems have been applied to describe the spatial organization of a broad array of cellular processes[31], ranging from protein oscillations in *E coli*[32], to coordination of mitotic signalling[33], to pattern formation in developing embryos[34, 35]. The response of actively moving particles to spatially heterogeneous, diffusive regulators has also been extensively investigated in the context of chemotaxis[36]. In contrast to most chemotactic cells, however, mitochondria have no currently known mechanism for directly sensing glucose gradients. Instead, they are expected to accumulate in response to local glucose concentration only. Our goal is to delineate the regimes in which such a crude form of chemotaxis can lead to substantial spatial organization and enhancement of metabolism.

Specifically, we model the modulation of mitochondrial density with glucose concentration in a tubular axonal region, focusing on two forms of spatial heterogeneity. In one case, we consider an axonal domain between two localized regions of glucose entry, representing the internodal region between nodes of Ranvier in myelinated neurons (Fig. 1a). The second case focuses on an unmyelinated cellular region with continuous glucose permeability, embedded in an external glucose gradient (Fig. 1b). In both cases, we show that mitochondrial accumulation and substantially enhanced metabolic flux is expected to occur over a limited range of glucose concentrations, which overlaps with physiological brain glucose levels. Our simplified quantitative model allows identification of a handful of key parameters that govern the extent to which glucose-mediated mitochondrial halting can modulate metabolism. We establish the region of parameter space where this mechanism has a substantial effect, and highlight its potential importance in neuronal metabolic flexibility and ability to respond to spatially varying glucose.

**FIG. 1.**
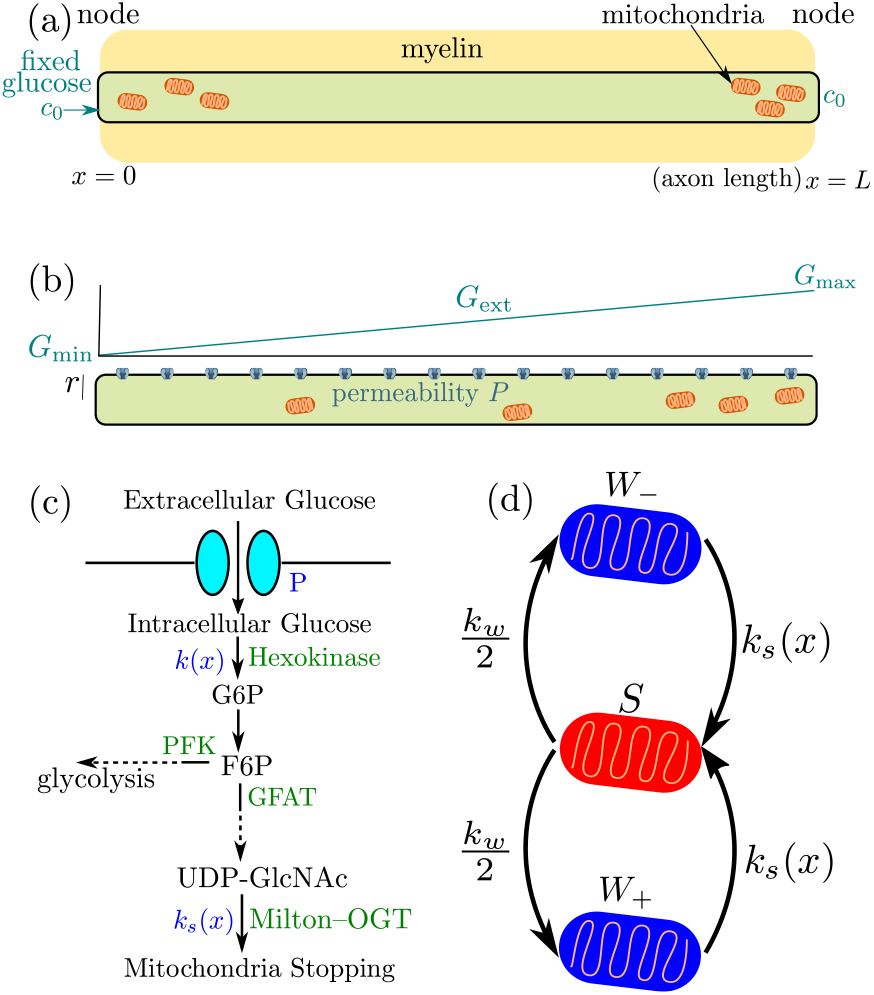
Schematic diagram of our simplified model for glucose-mediated mitochondrial transport regulation. (a) Myelinated axonal region, with glucose entry localized at the nodes of Ranvier. Mitochondria accumulate at nodes due to the higher glucose concentration (b) Unmyelinated axonal region, subject to a linear glucose gradient. Glucose permeability is uniform throughout, with mitochondrial accumulation occuring at the region of high external glucose (c) Key steps of the metabolic pathway linking glucose availability and mitochondrial halting. (d) Mitochondrial transport states and rates of transition between them (*W±* represents retrograde and anterograde motion, *S* represents the stationary state).

## MINIMAL MODEL FOR MITOCHONDRIAL AND GLUCOSE DYNAMICS

We begin by formulating a quantitative model to describe the spatial localization of mitochondria that halt in a glucose-dependent manner, in the presence of localized sources of glucose. This situation arises in myelinated neurons, which have glucose transporters enriched at the nodes of Ranvier, leading to highly localized sources of glucose spaced hundreds of micrometers apart within the cell[37].

Neuronal glucose transporters are known to be bidirectional[38], allowing glucose concentration within the cell to equilibrate with external glucose. For simplicity, we assume rapid transport of glucose through these transporters, so that the internal concentration of glucose at the nodes where transporters are present is assumed to be fixed. The cellular region between two glucose sources is modeled as a one-dimensional interval of length *L* with glucose concentration fixed to a value *c*_0_ at the interval boundaries (Fig.1a). Glucose diffuses throughout this interval with diffusivity *D*, while being metabolized by hexokinase enzyme in the first step of mammalian glucose utilization (Fig. 1c) [39].

The concentration of glucose is thus governed by the reaction-diffusion equation,

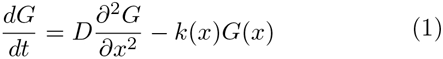

where *k*(*x*) describes the spatial distribution of the hexokinase enzyme as well as the rate of consumption. In the case of spatially uniform, linear consumption [*k*(*x*) = *k*, a constant] this equation can be solved directly, yielding a constant] this equation can be solved directly, yielding a distribution of glucose that falls exponentially from each source boundary, with a decay length 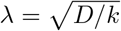 [40].

Hexokinase 1 (HK1), the predominant form of hexokinase expressed in neurons, is known to localize preferentially to mitochondria[41], which in mammalian axons can form individual organelles approximately 1*μ*m in length[42]. We carry out numerical simulations of Eq. 1 where consumption is limited to locations of individual discrete mitochondria, represented by short intervals of length ∆. Specifically, we define the mitochondria density as *M* (*x*) = *n*(*x*)*/*(*πr*^2^∆), where *n*(*x*) is the number of mitochondria overlapping position *x*, and *r* is the axon radius. The phosphorylation of glucose by mitochondrial hexokinase is assumed to follow Michaelis-Menten kinetics, described by

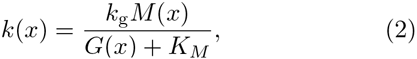

where *K_M_* is the saturation constant and *k*_g_ is the turnover rate of glucose (per unit time per mitochondrion). The turnover rate *k_g_* incorporates both the catalytic rate of hexokinase and the number of hexokinase enzymes per mitochondrion. This expression reduces to the case of constant linear consumption when glucose concentration is low (*G* ⪡ *K_M_*) and mitochondria are uniformly distributed throughout the region.

In general, glucose consumption depends on the location of mitochondria within the domain. Mitochondrial distribution in neurons is known to be mediated through regulation of their motor-driven motility[24, 28]. Individual mitochondria switch between processively moving and paused states, modulated by the interplay between kinesin and dynein motors and the adaptor proteins that link these motors to the mitochondria[43]. In our model, we simulate mitochondria as stochastically switching between a processive walking state that moves in either direction with velocity *v* and a stationary state. The rate of initiating a walk (*k_w_*) is assumed to be constant, while the halting rate (*k_s_*(*x*)) can be spatially heterogeneous. For simplicity, we assume the mitochondria are equally likely to move in the positive (+) or negative (−) direction each time they initiate a processive walk (Fig. 1b). We note that motile axonal mitochondria tend to move consistently in an anterograde or retrograde manner, with sporadic pauses followed by motion in the same direction[26, 44]. While such dynamics will modify the time-scale for establishing a mitochondrial distribution in response to glucose heterogeneity, the stationary state distribution depends only on the net fraction of mitochondria moving in each direction. Here, we focus on these stationary distributions, while assuming equal frequency of anterograde and retrograde transport.

It has recently been demonstrated that a key motor adaptor protein (Milton) is sensitive to glucose levels, halting mitochondrial motility when it is modified through O-GlcNAcylation by the OGT enzyme[28]. Upon entry into the cell, the first rate-limiting step of glucose metabolism is its conversion into glucose-6-phosphate by hexokinase. Further downstream metabolic pathways split, with much of the flux going to glycolysis while a small fraction is funneled into the hexosamine biosynthetic pathway (HBP). This pathway produces UDP-GlcNAc, the sugar substrate for O-GlcNAcylation (Fig. 1c)[45]. In our model, we assume that the rate of UDP-GlcNAc production equals the rate of glucose conversion by hexokinase, scaled by the fraction of G6P that is channeled into the hexosamine pathway. This assumption is valid if the saturation constants for the first pathway step separating glycolysis and hexosamine biosynthesis are comparable to each other (see Supplementary Information). This in fact appears to be the case for glutamine–fructose-6-phosphate aminotransferase (GFAT, *K_M_* ≈ 1mM[46]), which initiates the hexosamine pathway, and phosphofructokinase (PFK, *K_M_* ≈ 1.3mM[46]), which funnels intermediates to the glycolytic pathway. In this case, saturation of the initial glucose conversion step will imply saturation of the entire hexosamine biosynthetic pathway. We therefore model the kinetics of Milton modification using the same Michaelis-Menten form as for hexokinase activity, with the pathway flux leading to Milton modification subsumed within a rate constant for mitochondrial stopping (*k_s_*).

We note that the subcellular organization of the intermediates in the conversion from glucose into O-GlcNAcylated Milton is largely unknown. In our model, we make the extreme case assumption that all intermediates are localized to mitochondria, with only the initial glucose substrate capable of diffusing through the cytoplasm. We note that cytoplasmic diffusion of any of the pathway intermediates would attenuate the effect on mitochondrial localization. Our simplified model thus gives an upper limit on the extent to which mitochondria can localize at high glucose regions through the Milton modification mechanism. Following these simplified assumptions, we treat the kinetics of mitochondrial halting as dependent only on the local glucose concentration, according to the functional form

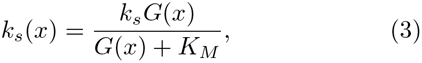

where *K_M_* is the Michaelis-Menten constant of hexokinase.

We proceed to evolve the simulation forward in time, with glucose consumption localized to regions within ±∆/2 Supplementary Information). A snapshot of one simulation run is shown in Fig. 2a, highlighting the accumulation of stationary mitochondria in the high glucose regions near the ends of the domain.

**FIG. 2.**
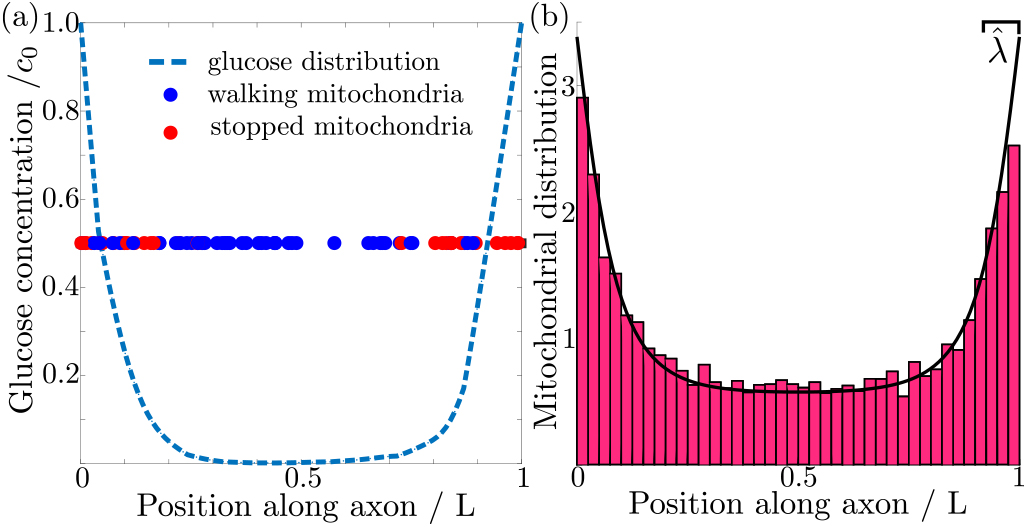
(a) Glucose distribution and position of individual mitochondria (b) Normalized itochondrial distribution, 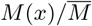, obtained from simulating discrete mitochondrial motion (histogram compiled from 100 independent simulations), compared to numerical calculation of steady state continuous mitochondrial disribution (black curve). Results shown are for parameter values: λ ̂ = 0.08, *c*̂_0_ = 1, *k*̂_*s*_ = 100.

We are interested primarily in investigating the steady-state distribution of mitochondria and glucose in this system, averaged over all possible mitochondrial trajectories. We thus proceed to coarse-grain our model by treating the distribution of mitochondria as a continuous field *M* (*x*) = *W*_+_(*x*) + *W_−_*(*x*) + *S*(*x*), where *W*_+_(*x*) is the distribution of mitochondria walking in the positive direction, *W_−_*(*x*) is the distribution of those walking in the negative direction, and *S*(*x*) is the distribution of stationary mitochondria. We can then write down the coupled differential equations governing the behavior of the mitochondrial distributions as:

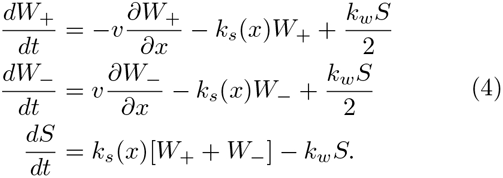

The glucose distribution evolves according to Eq. 1 with consumption rate *k*(*x*) given by Eq. 2. The boundary conditions at the ends of the domain are assumed to be reflective for the mitochondrial distributions, and to have a fixed glucose concentration *c*_0_. The stationary state for this system can be calculated numerically (see Supplementary Information). The formulation with a continuous mitochondrial density faithfully represents the behavior of simulations with discrete mitochondria, as illustrated in Fig. 2b.

The steady-state spatial distribution of mitochondria and glucose in the continuous system depend on six parameters: *k_s_/k_w_, K_M_, c*_0_*, D, L, k_g_M*̅ where *M̅* is the average mitochondrial density in the axon (number of mitochondria per unit volume). Estimates of physiologically relevant values are provided in Table I. Dimensional analysis indicates that three of these parameters can be used to define units of time, length, and glucose concentration, leaving three dimensionless groups. We choose to use the following three dimensionless parameters, each of which has an intuitive physical meaning:

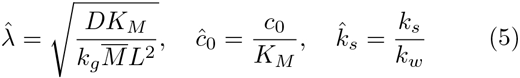

**TABLE I.**
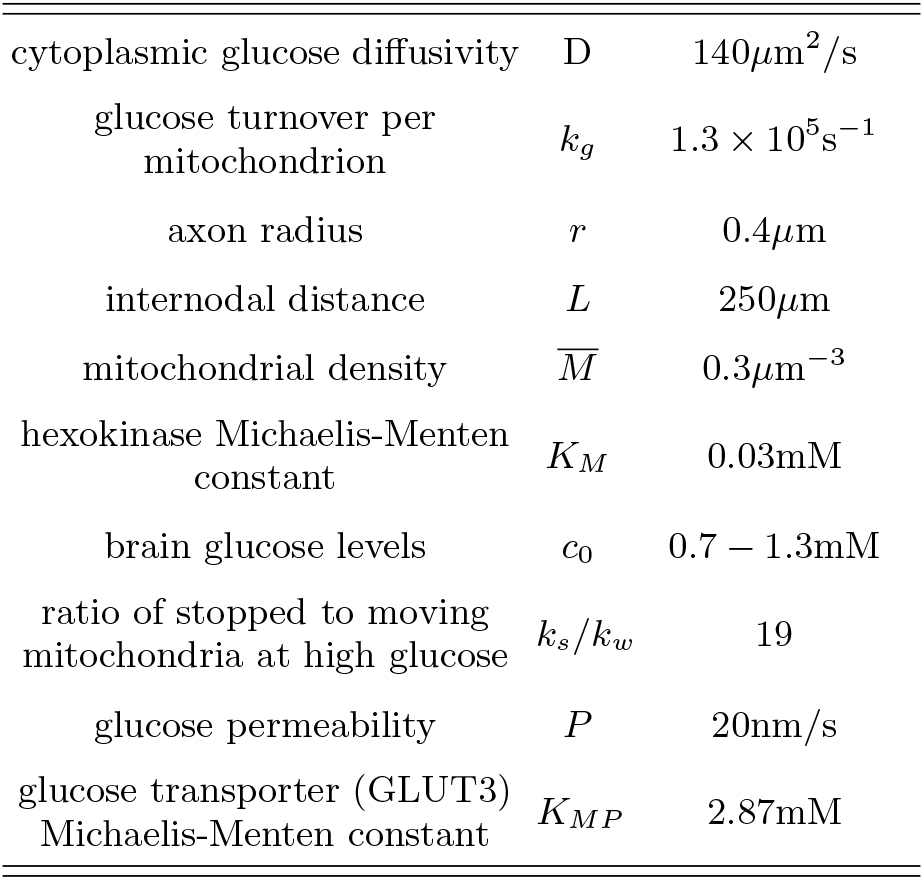
Physiological parameter values estimated from published data. Details of estimates are provided in the Supplementary Information.

Here λ̂ is the length-scale of glucose decay relative to the domain length, *c*̂_0_ is the boundary glucose concentration relative to the saturation constant *K_M_*, and *k*̂_*s*_ is the ratio of stopped to walking mitochondria at high glucose levels. We proceed to explore the steady-state distribution of mitochondria and glucose as a function of these three parameters.

## MITOCHONDRIAL LOCALIZATION REQUIRES LIMITED RANGE OF EXTERNAL GLUCOSE

In order for mitochondria to preferentially accumulate at the source of glucose via a glucose-dependent stopping mechanism, three criteria must be met. First, the glucose concentration needs to be higher at the source than in the bulk of the cell, as occurs when the decay length due to consumption is much smaller than the size of the domain (λ ̂ ≪ 1). Second, if glucose levels become too high (*c*_0_ ̂ ≫ 1) then both glucose consumption rates and stopping rates of the mitochondria become saturated, leading to a flattening of glucose and mitochondrial distributions (Fig. 3). There is thus an upper limit on the possible external glucose concentrations that will yield mitochondrial localization at the edges of the domain. Finally, the mitochondria must spend a substantial amount of time in the stationary state, since walking mitochondria will be broadly distributed throughout the domain. Because the stopping rate is itself dependent on the glucose concentration, this criterion implies that very low concentrations will also not allow mitochondrial localization. Fig. 3 shows the distribution of glucose and mitochondria at different values of the external glucose *c*̂_0_, illustrating that accumulation of mitochondria at the edges requires intermediate glucose levels.

**FIG. 3.**
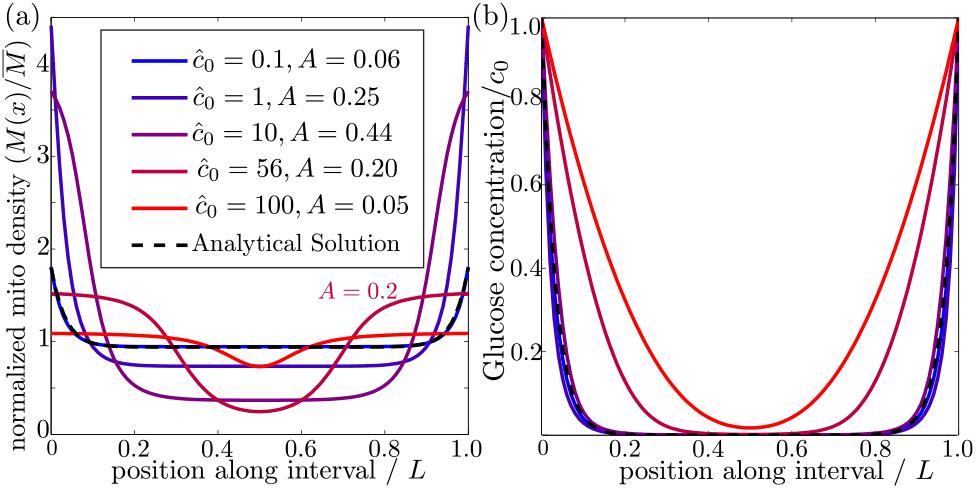
Effect of external glucose concentration on intracellular glucose and mitochondrial dist<u>rib</u>utions. (a) Normalized mitochondrial distribution (*M* (*x*)*/M* ̅), for different values of edge concentration *c*̂_0_. The curve with *c*̂0 = 56 illustrates the accumulation cutoff *A* = 0.2. (b) Glucose distribution normalized by edge concentration (*G*(*x*)*/c*_0_). The black dashed line in both panels indicates the analytical solution for the low glucose limit.

To characterize the distribution of mitochondria along the interval, we introduce an accumulation metric *A*, defined by

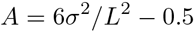

where *σ*^2^ is the variance in the mitochondrial distribution. This metric scales from *A* = 0 for a uniform distribution to *A* = 1 for two narrow peaks at the domain edges. Mitochondrial distributions with several different values of the accumulation metric are shown in Fig. 3a. We use a cutoff of *A* = 0.2 to define distributions where the mitochondria are localized at the glucose source.

We explore the dependence of the mitochondrial accumulation on the three dimensionless parameters defining the behavior of the system: the stopping rate constant *k*̂_*s*_, the glucose decay length λ̂, and the external concentration *c*̂_0_. Because only the stopped mitochondria localize near the glucose sources, increasing the fraction of mitochondria in the stopped state (increased *k*̂_*s*_) inevitably raises the overall accumulation (Fig. 4a). The fraction of mitochondria in the stopped state will depend on both *k*̂_*s*_ and the overall levels of glucose, as dictated by *ĉ*_0_ (Fig. 4b). Experimental measurements indicate that at high glucose concentrations, approximately 95% of mitochondria are in the stationary state[28]. We are thus interested primarily in the parameter regime of high stopping rates: *k*̂ ≳_*s*_ 10. The limited range of concentrations that lead to mitochondrial accumulation at the edges of the domain can be seen in Fig. 4a.

**FIG. 4.**
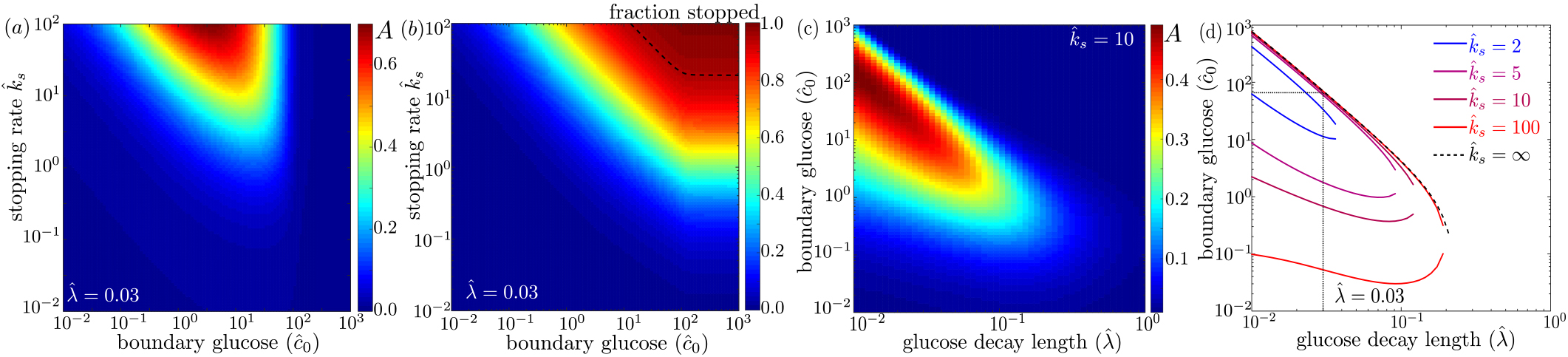
Effect of model parameters on mitochondrial accumulation at regions of localized glucose entry. (a) Accumulation metric as a function of boundary glucose levels and mitochondrial stopping rate. (b) Fraction of mitochondria in the stopped state. Black dashed line indicates parameters corresponding to 95% stopped mitochondria. (c) Accumulation metric as a function of glucose levels *c*̂_0_ and decay length λ̂. (d) Phase diagram for mitochondrial accumulation, showing upper and lower concentration cutoffs for accumulation above the cutoff of *A*_cut_ = 0.2. Dashed black line shows limit of high stopping rate *k*̂*s*. Dotted black line indicates estimate of λ̂ for physiological parameters, and corresponding upper concentration cutoff.

For a high stopping rate (*k*̂_*s*_ = 10), we then calculate the mitochondrial accumulation as a function of the remaining two parameters: λ̂*, c*̂_0_. Here, again, we note that only intermediate glucose concentrations result in accumulation, with the range of concentrations becoming narrower as the decay length λ̂ becomes comparable to the domain size (Fig. 4c). We can establish the concentration range within which substantial accumulation is expected, by setting a cutoff *A* = 0.2 on the accumulation metric and calculating the resulting phase diagram (Fig. 4d). Below the lower concentration cutoff, insufficient mitochondria are in the stationary state and so no localization is seen. This lower cutoff disappears in the limit of infinite *k*̂_*s*_. At intermediate concentrations, mitochondria are localized near the domain edges. Above the upper concentration cutoff, no localization is observed due to saturation of the Michaelis-Menten kinetics.

Using empirically derived approximations for the rate of glucose consumption by mitochondria and the diffusivity of glucose in cytoplasm (see Table I), we estimate the decay length parameter as λ̂ ≈ 0.03. The mitochondria are then expected to localize near the glucose source only if *c*̂_0_ *<* 66. Because the saturation concentration for hexokinase is quite low (*K_M_ ≈* 0.03mM)[39], we would expect mitochondrial accumulation for glucose concentrations below about 2 mM. We note that physiological brain glucose levels have been measured at 0.7 – 1.3mM, depending on the brain region[47], implying that glucose-dependent halting of mitochondrial transport would be expected to result in localization of mitochondria at nodes of Ranvier.

## GLUCOSE-DEPENDENT HALTING CAN INCREASE METABOLIC FLUX UNDER PHYSIOLOGICAL CONDITIONS

Localizing mitochondria to the glucose entry points is expected to increase the flux of glucose entering the cell, thereby potentially enhancing the overall metabolic rate. We calculate the overall effect of transport-based regulation on the net metabolic flux within the simplified model with localized glucose entry. Fig. 5 shows the effect of increasing mitochondrial stopping rates (*k*̂_*s*_) on the total rate of glucose consumption in the interval between nodes of glucose influx. At low *k*̂_*s*_ values, mitochondria are distributed uniformly throughout the interval. At high *k*̂_*s*_ values and at sufficiently low glucose concentrations, the mitochondria cluster in the regions of glucose entry, increasing the overall consumption rate by up to 40% at physiologically relevant glucose levels (*c*_0_ = 1mM). We note that in hypoglycemic conditions, glucose levels can drop to 0.1mM [48], further increasing the magnitude of this effect.

**FIG. 5.**
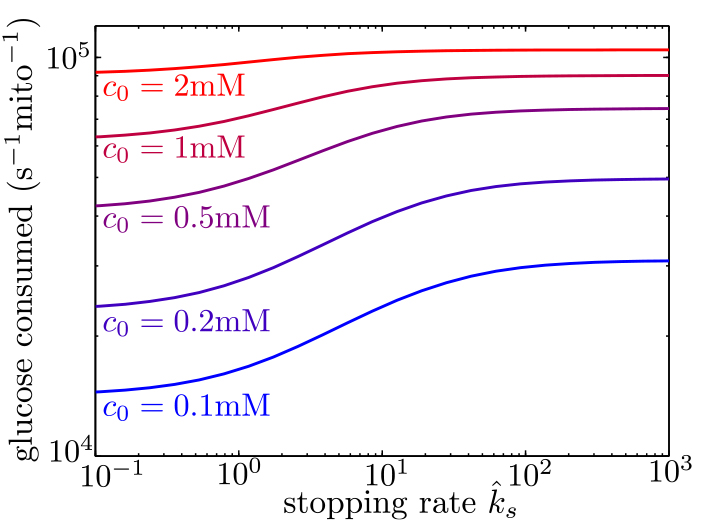
Mitochondrial stopping increases overall metabolic flux. Total glucose consumption per mitochondrion, averaged over the full interval, is shown for different edge glucose concentrations (*c*_0_) as a function of the mitochondrial stopping rate *k*̂_*s*_. The limit of small *k*̂_*s*_ corresponds to uniform mitochondria distribution. Parameters for the model are taken from Table I.

In the case of limited glucose transport into the cell, intracellular glucose levels could be significantly below the concentrations outside the cell. Measurements of intracellular glucose in a variety of cultured mammalian cell types indicate internal concentrations within the range of 0.07 – 1mM, up to an order of magnitude lower than glucose concentrations in the medium[49]. However, neuronal cells are known to express a particularly efficient glucose transporter (GLUT3)[50], and these transporters have been shown to be highly concentrated near the nodes of Ranvier[12, 14]. We therefore assume that glucose import into the nodes is not rate limiting for myelinated neurons in physiological conditions. Introducing a finite rate of glucose transport would effectively decrease the intracellular glucose concentration at the nodes *c*_0_, increasing the enhancement in metabolic flux due to mitochondrial localization. In subsequent sections, we explore the role of limited glucose import in unmyelinated axons with spatially uniform glucose permeability.

## MODEL FOR SPATIAL ORGANIZATION IN A GLUCOSE GRADIENT

Extracellular brain glucose levels exhibit substantial regional variation, particularly under hypoglycemic conditions where more than ten-fold differences in local glucose concentrations have been reported[51]. Because individual neurons can traverse multiple different brain regions[19], a single axon can be subjected to heterogeneous glucose levels along its length. This raises the possibility that glucose-dependent mitochondrial localization can play a role in neuronal metabolic flexibility even in the case where glucose entry into the cell is not localized to distinct nodes. We thus extend our model to quantify the distribution of mitochondria in an axon with limited but spatially uniform glucose permeability that is subjected to a gradient of external glucose. This situation is relevant, for instance, to unmyelinated neurons in infant brains, as well as to *in vitro* experiments with neurons cultured in a glucose gradient[28].

In this model, the extracellular environment provides a continuous source of glucose whose influx is limited by the permeability of the cell membrane. Intracellular glucose dynamics are then defined by the reaction-diffusion equation

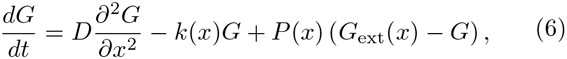

where the first term corresponds to diffusive glucose spread, the second to a spatially varying metabolism of glucose, and the third to the entry of glucose into the cell. Here, *G*_ext_ is the external glucose concentration, and *P* (*x*) is the membrane permeability to glucose, which we assume to depend in a Michaelis-Menten fashion on the difference between external and internal glucose concentration:

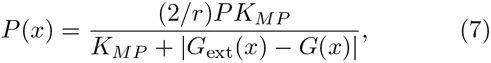

where *P* is the spatially uniform permeability constant in units of length per time. This functional form incorporates two known features of glucose transporters: (1) they are bidirectional, so that the overall flux through the transporter at low glucose levels should scale linearly with the difference between external and internal glucose[52]; (2) neuronal glucose transporters saturate at high glucose levels (GLUT3 *K_MP_ ≈* 2.87mM[53], with an even higher saturation constant for GLUT4 [54]). When the difference in glucose levels is low, the overall flux of glucose entering the cell reduces to *P* (*G*_ext_(*x*) – *G*(*x*)). Mitochondria dynamics are defined as before (Eq. 4), and we again assume Michaelis-Menten kinetics for glucose metabolism by hexokinase localized to mitochondria (Eq. 2).

We note that the dynamics in Eq. 6 are governed by three time-scales: the rate of glucose transport down the length of the axon, rate of glucose consumption, and rate of glucose entry. The first of these rates becomes negligibly small in the limit 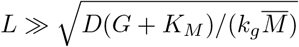. Because internal glucose levels can never exceed the external concentrations, in the range where *G*_ext_ < 10mM, the rate of diffusive transport should become negligible for *L* ≫ 150*μ*m. In the limit where intracellular glucose is much less than *K_m_*, this criterion reduces to λ̂ ≪ 1, indicating that glucose diffuses over a very small fraction of the interval before being consumed. The interval length *L* in this model represents an axonal length which can range over many orders of magnitude. We focus on axon lengths above several hundred microns, allowing us to neglect the diffusive transport of intracellular glucose (see Supplementary Information, Fig. S1).

The steady-state glucose profile can then be determined entirely by the local concentration of mitochondria and external glucose. For a given mitochondrial density *M* (*x*) and external glucose profile *G*_ext_(*x*), the corresponding intracellular glucose concentration can be found directly by solving the quadratic steady-state version of Eq.6 without the diffusive term. However, the steady-state mitochondrial distribution cannot be solved locally, because the limited number of mitochondria within the axon couples the mitochondrial density at different positions. We thus employ an iterative approach to numerically compute the steady-state solution for both glucose and mitochondrial density under a linear external glucose gradient 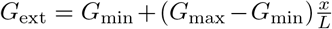 (see Supplementary Information).

For parameter combinations where intracellular glucose concentrations are above *K_M_* but well below *G*_ext_, the entry and consumption processes for glucose are both saturated. There is then a steep transition between two different regimes. In one regime, glucose entry exceeds consumption and internal glucose levels approach the external concentrations. In the other, consumption dominates and glucose levels drop below saturating concentrations. The key dimensionless parameter governing this transition can be defined as the ratio of entry to consumption rates:

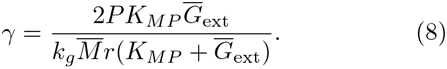

This ratio can be modulated in the cell either by recruiting varying amounts of glucose transporters (adjusting *P*) or changing the total amount of active hexokinase (adjusting *k_g_M ̅*).

The remaining dimensionless parameters determining the behavior of this simplified model are the external glucose concentration relative to the hexokinase saturation constant (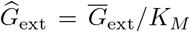), the relative magnitude of the glucose gradient, 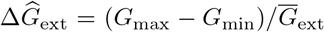, the ratio of stopped to walking mitochondria *k̂_s_* = *k_s_/k_w_*, and the saturation constant for glucose transporters *K_MP_ /K_M_ ≈* 96. The last parameter is expected to remain approximately constant in neuronal cells. The average external glucose concentration and glucose gradient are expected to vary substantially depending on the glycemic environment to which the neuron is exposed. We note that 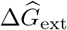 has a maximum possible value since the minimal glucose concentration cannot drop below 0. We proceed to analyze the limiting case where the glucose gradient is as steep as possible for any given value of average external glucose (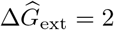).

## MITOCHONDRIAL ARREST ENABLES METABOLIC ENHANCEMENT UNDER GLUCOSE GRADIENT

We quantify the amount of mitochondrial accumulation at the high glucose side of the domain by calculating the total mitochondrial density within the distal 10% of the interval compared to a uniform distribution, in analogy to experimental measurements[28]. Substantial enrichment in the high glucose region occurs when glucose entry into the cell cannot keep up with consumption (γ ≪ 1) and the intracellular glucose levels drop below the hexokinase saturation concentration *K_M_*, as can be seen in the glucose and mitochondrial distributions plotted in Fig. 6a-c. The interplay between external glucose levels and the entry / consumption rates is illustrated in Fig. 6d. For external glucose concentrations well above *K_M_* there is a sharp transition to mitochondrial enrichment at γ *<* 1. At the lowest levels of intracellular glucose, accumulation is again reduced because a very small fraction of mitochondria are found in the stopped state. In the limit of high *k_s_*, mitochondrial accumulation would occur for arbitrarily low values of γ (Fig. S2). We note that because glucose entry and turnover are much faster than diffusive spread for biologically relevant parameter regimes, the model results do not depend on the cell length *L* (Supplementary Information and Fig. S1).

**FIG. 6.**
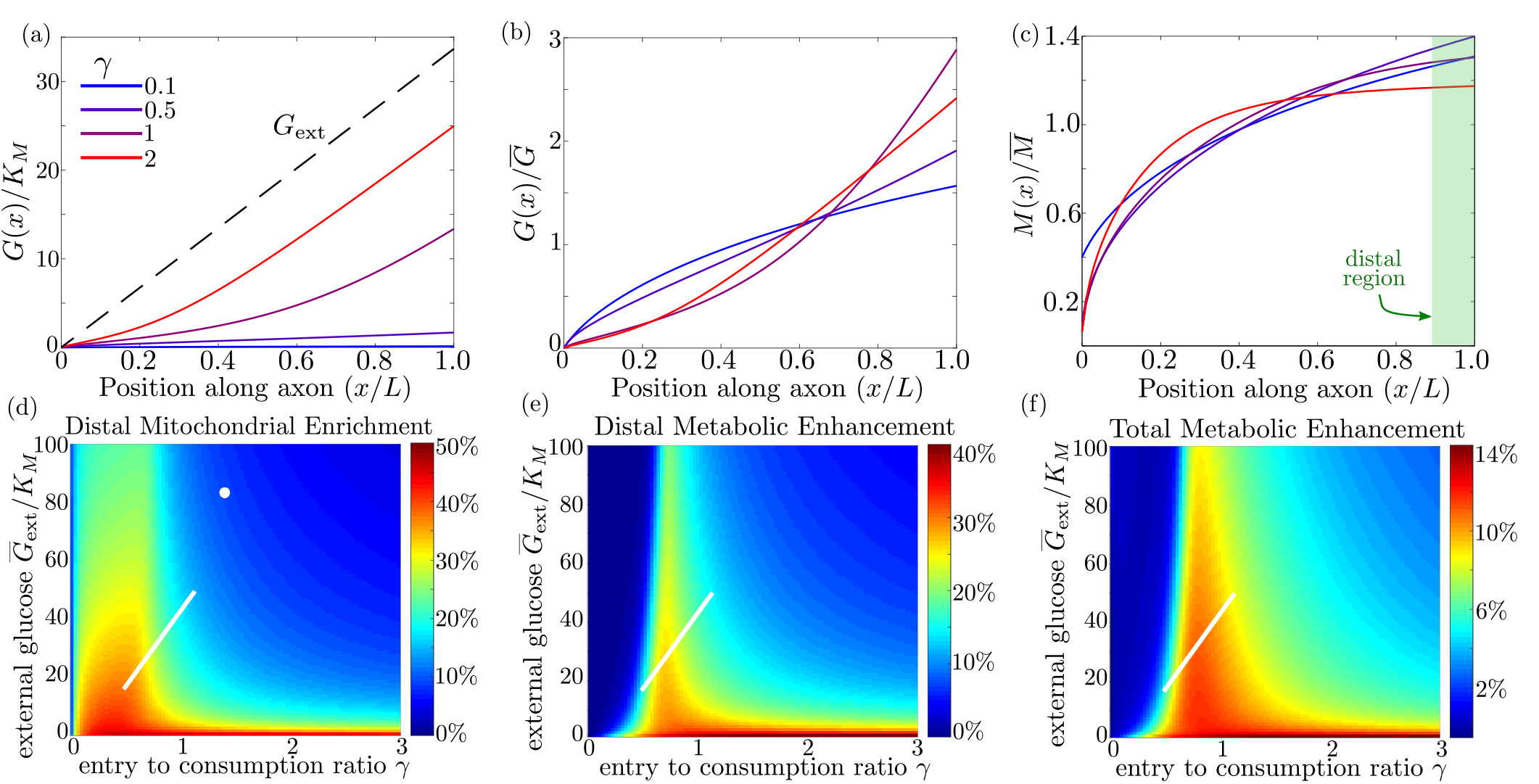
Mitochondrial and glucose organization in a region with uniform glucose permeability, subjected to a gradient of external glucose. (a) Internal glucose levels for the steady state solution with *G*̅_ext_*/K_M_* = 17 (*G*̅_ext_ = 0.5 mM) and varying ratios of entry to consumption rate γ. Black dashed line shows external glucose levels. (b) Corresponding normalized distribution of internal glucose. (c) Corresponding normalized mitochondrial distribution. Shaded box indicates distal region used for calculating mitochondrial enrichment and metabolic enhancement in panels d-e. (d) Mitochondrial enrichment in the distal 10% of the interval at highest external glucose, compared to a uniform distribution. White dot marks estimated parameter values for neuronal cell culture experiments (*G*̅_ext_ = 2.5mM). (e) Enhancement in metabolic flux in the distal region at high glucose, compared to a uniform mitochondrial distribution. (f) Enhancement in metabolic flux over full interval. White line in (d-f) shows estimated parameter range for physiological glycemic levels 0.5mM *< G*̅_ext_ *<* 1.5mM. Parameter values *k*̂*s* = 19, 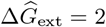 used throughout.

Experimental measurements of mitochondrial enrichment in cultured neurons subjected to a gradient of 0 to 5mM glucose have indicated an approximately 20% enrichment in mitchondrial counts at the axonal region exposed to high glucose. We note that using published estimates of typical glucose permeability and mitochondrial glucose turnover for mammalian cells (Table I) yields a ratio of entrance and consumption rates of γ ≈ 1.9 for this experimental system. Because this ratio is above 1, we would not expect to see substantial mitochondrial enrichment. To result in the experimentally observed enrichment at high glucose, the ratio γ would need to be reduced by approximately a factor of 2, implying the existence of additional regulatory mechanisms. Modulation of γ could be achieved by either decreasing the number of glucose transporters in the cell (reducing *P*) or upregulating total hexokinase levels (increasing *k_g_*). Neurons are believed to regulate both the density of glucose transporters and hexokinase activity in response to external glucose concentrations and varying metabolic demand[55–57]. In particular, adaptation to glycemic levels well above physiological values, as well as possibly reduced synaptic activity in a cultured environment, may result in downregulation of glucose transporters, lowering the value of γ. The discrepancy between model prediction and observed mitochondrial accumulation highlights the existence of additional regulatory pathways not included in the current model whose role could be explored in further studies that directly quantify glucose entry and consumption rates in cultured neurons.

Physiological brain glucose levels have been measured at 0.7mM - 1.3mM[47], with hypoglycemic levels dipping as low as 0.1mM and hyperglycemic levels rising up to 4mM[48]. Axons that stretch across different brain regions with varying glucose levels can thus be subject to a glucose gradient with *G̅*_ext_ on the order of 1mM (white line on Fig. 6d). We note that the physiological range overlaps substantially with the region of high mitochondrial accumulation, indicating that glucose-dependent halting can modulate mitochondrial distribution under physiologically relevant glycemic levels.

By accumulating mitochondria at the cellular region subjected to higher external glucose, the metabolic flux in that region can be substantially enhanced. In Fig. 6e we plot the enhancement in glucose consumption rates (compared to the case with uniformly distributed mitochondria) within the 10% of cellular length subjected to the highest glucose concentrations. Metabolic enhancement occurs within a narrow band of the γ parameter. The drop-off in enhancement at low values of the internal glucose concentration (low γ) is due to the coupling between glucose levels and mitochondrial localization. Specifically, mitochondrial accumulation at the region subject to high glucose concentration increases the local rate of consumption in that region, driving down local internal glucose levels. Consequently, the difference in internal glucose concentrations between the two ends of the cell is decreased when internal levels fall substantially below *K_M_* (Fig. 6b), reducing the enhancement of metabolic flux. Although mitochondrial accumulation decreases metabolic flux in the low glucose region, the total rate of glucose consumption integrated throughout the cell is enhanced by up to approximately 14% when γ ≈ 1 (Fig. 6f).

It is interesting to note that the typical physiological range of external glucose levels spans the narrow band of parameter space where metabolic enhancement is expected (white lines on Fig. 6e,f). These results implicate glucose-dependent mitochondrial stopping as a quantitatively plausible mechanism of metabolic flexibility, increasing metabolism in regions with high nutrient availability for axonal projections that span between hypoglycemic and euglycemic regions. The magnitude of this effect can be tightly controlled by the cell through modulating overall rates of glucose entry and consumption. Thus, by coupling mitochondrial transport to local glucose levels, whole-cell changes in hexokinase or glucose transporter recruitment can be harnessed to tune the cell’s response to spatially heterogeneous glucose concentrations.

## DISCUSSION

The minimal model described here provides a quantitative framework to explore the interdependence of glucose levels and mitochondrial motility and their combined effect on neuronal metabolic flux. Glucose-mediated halting of mitochondrial transport is shown to be a plausible regulatory mechanism for enhancing metabolism in cases with spatially heterogeneous glucose availability in the neuron.

We have quantitatively delineated the regions in parameter space where such a mechanism can have a substantial effect on mitochondrial localization and metabolic flux. Specifically, mitochondrial positioning requires both sufficient spatial variation in intracellular glucose and sufficiently low absolute glucose levels compared to the saturation constant of the hexokinase enzyme. In the case of tightly localized glucose entry (as at the nodes of Ranvier), intracellular spatial heterogeneity requires a small value of the dimensionless length scale for glucose decay (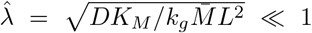). For physiologically estimated values, mitochondrial localization to the nodes is expected to occur for glucose levels below approximately 2mM, comparable to physiological brain glucose concentrations[47, 49]. In the case where glucose can enter homogeneously throughout the cell surface (as with unmyelinated axons), heterogeneity can arise from an external glucose gradient. We show that metabolic enhancement through mitochondrial positioning occurs in a narrow range of the key parameter γ = (2*PK_MP_G̅*_ext_)*/*(*k_g_M̅*(*K_MP_* + *G̅*_ext_)), which describes the ratio of glucose entry to glucose metabolism, and that this narrow range intersects with physiological estimates.

The model developed here is intentionally highly simplified, encompassing a minimal set of parameters necessary to describe glucose-dependent mitochondrial localization. Other regulatory pathways that determine mitochondrial positioning are not included in this basal model. In particular, we do not consider here calcium-based transport regulation, which is known to localize mitochondria to regions of synaptic activity[26, 27, 29, 58]. Upregulating OGT signaling in cultured cells has been shown to decrease the fraction of motile mitochondria by a factor of three, while reducing endogenous OGT nearly doubles the motile fraction, indicating that a substantial number of stationary mitochondria are stopped as a result of OGT activity[28]. Our model assumes the extreme case where all stopping events are triggered in a glucose-dependent manner, thereby isolating the effect of glucose heterogeneity. Stopping mechanisms dependent on neuronal firing activity could alter mitochondrial distribution in concert with glucose-dependent halting, increasing the density of mitochondria at presynaptic boutons or near areas of localized calcium influx as at the nodes of Ranvier[58]. We note that mitochondria have previously been shown to accumulate at spinal nodes of Ranvier in response to neuronal firing activity[58, 59]. The mechanism described here provides an additional driving force for mitochondrial localization near the nodes even in quiescent neurons.

Several key parameters that regulate mitochondrial localization in response to glucose heterogeneity can be dynamically regulated in neurons. Specifically, the rate of glucose consumption (*k_g_M̅*) can be tuned by modulating the concentration or activity of hexokinase within mitochondria or by altering total mitochondrial size and number. This parameter controls both the glucose decay length λ̂ in the case of localized glucose influx and the ratio of glucose entry to consumption γ in the case of spatially distributed entry. We note that our model assumes hexokinase to be localized exclusively to mitochondria. The predominant form of hexokinase in the brain (HK1) is known to bind reversibly to the mitochondrial membrane, with exchange between a mitochondria-bound and a cytoplasmic state believed to contribute to the regulation of its activity[60]. Release of hexokinase into the cytoplasm would result in more spatially uniform glucose consumption, negating the metabolic enhancement achieved through mitochondrial localization.

An additional parameter known to be under regulatory control is the rate of glucose entry into the neuron (*P*). The glucose transporters GLUT3[9, 50, 57] and GLUT4[17] have been shown to be recruited to the plasma membrane in response to neuronal firing activity. Interestingly, transporter densities are themselves spatially heterogeneous, concentrating near regions of synaptic activity[17, 61]. The model described in this work quantifies the extent to which a locally increased glucose influx can enhance total metabolic flux, given the ability of mitochondria to accumulate at regions of high intra-cellular glucose.

A number of possible feedback pathways linking glucose distribution and mitochondrial positioning are not included in our basic model. For instance, hexokinase release from mitochondria into the cytoplasm (potentially altering *k_g_*) is known to be triggered at least in part by glucose-6-phosphate, the first byproduct in glucose metabolism[62]. Chronic hypoglycemia has been linked to an upregulation in GLUT3 in rat neurons [63], which would in turn lead to an increased glucose uptake (*P*). The fraction of glucose funneled into the hexosamine biosynthetic pathway (incorporated within *k_s_*) can also be modified through feedback inhibition of GFAT by the downstream metabolic product UDP-GlcNAc[46]. Such feedback loops imply that several of our model parameters (*P*, *k_g_*, *k_s_*) are themselves glucose-dependent and may become spatially non-uniform in response to heterogeneous glucose. Incorporating these effects into a spatially resolved model of metabolism would require quantifying the dynamics of both the feedback pathways and mitochondrial positioning, and forms a promising avenue for future study.

Control of glucose entry and consumption underlies cellular metabolic flexiblity, and defects in the associated regulatory pathways can have grave consequences for neuronal health. Misregulation of hexokinase has been highlighted as a contributor to several neurological disorders, ranging from depression [64] to schizophrenia [65]. Neuronal glucose transporter deficiency has been linked to autism spectrum disorders[66] and Alzheimer’s disease[67]. Furthermore, defects in mitochondrial transport, with the consequent depletion of mitochondria in distal axonal regions, contribute to peripheral neuropathy disorders[22].

Glucose-dependent mitochondrial localization provides an additional layer of control, beyond conventionally studied regulatory mechanisms, which allows the cell to respond to spatial heterogeneity in glucose concentration. Our analysis paves the way for quantitative understanding of how flexible regulation of metabolism can be achieved by controlling the spatial distribution of glucose entry and consumption.

## AUTHOR CONTRIBUTIONS

All authors contributed to designing the research; AA and EFK developed analytical tools and wrote the manuscript; AA wrote and performed simulations and analyzed data; all authors read and edited the final manuscript.

## ACKNOWLEDGEMENTS

We thank Saurabh Mogre for fruitful discussions and Manho Tang for numerical modeling advice. This work was supposrted from the University of California San Diego, Chancellor’s Research Excellence Scholarship to A. A., from institutional funds to E. F. K. and G. P., and from NIH (R35GM128823) to G. P.

